# Imaging cerebral arteries tortuosity and velocities by transcranial Doppler ultrasound is a reliable assessment of brain aneurysm in mouse models

**DOI:** 10.1101/2022.01.14.476192

**Authors:** Héloïse Lebas, Alexandre Boutigny, Clémence Maupu, Jonas Salfati, Cyrille Orset, Mikael Mazighi, Philippe Bonnin, Yacine Boulaftali

## Abstract

**Background and Purpose:** Intracranial aneurysms (IAs) are common vascular abnormalities of the brain with a prevalence of 3.2% in the general population. In the past few decades, several pathophysiological processes leading to IA rupture were identified, including irregular IA shape, an altered hemodynamic stress within the IA and vessel wall inflammation. The use of preclinical models of IA and imaging tools are paramount to better understand the underlying disease mechanisms. Therefore, there is a need for imaging methods to monitor intracranial aneurysm formation.

**Methods:** We used two established mouse models of IA and we analyzed the progression of the IA by magnetic resonance imaging (MRI), transcranial Doppler (TCD), and histological studies.

**Results:** In both models of IA, we observed by TCD a significant decrease of the blood velocities and wall shear stress of the internal carotid arteries (ICA). We also observed the formation of tortuous arteries in both models which were correlated with the presence of an aneurysm as confirmed by MRI and histological analysis. A high grade of tortuosity has been associated with a significant decrease of the mean blood flow velocities and a greater artery dilation.

**Conclusions:** TCD is robust and easy imaging method to evaluate the progression of IA. The decrease of the blood flow velocities and the tortuosity can be used as reliable readout for IA detection.

## Introduction

Intracranial aneurysms (IA) are pathological focal dilatation of intracranial arteries mainly located on the circle of Willis. Their rupture, occurring in 6/100000 persons, has severe consequences including the death of 27 to 44% of patients ^1^. There are several identified risk factors that contribute to the formation of IA such as for example hypertension, smoking, family history of IA, head trauma or gender as women have an increased risk of IA formation^1^. In the past decade, several pathophysiological processes involved in the development and rupture of IAs were identified such as inflammation or an altered hemodynamic. Abnormal blood flow condition plays a key role in IA development explaining the IA location, commonly found at arterial bifurcations where excessive hemodynamic stresses are exerted on arterial walls ^2^. There is a close relation between wall shear stress, endothelial dysfunction and the downstream inflammatory reaction ^3^. The main hemodynamic parameter analysed is the wall shear stress (WSS) which is defined as the frictional force tangent to vessel wall induced by blood flow. High and low WSS have been determined to play a role in the formation and progression of IA ^4^. In particular, high shear stress has been reported to modify the local expression of several other genes that contribute to IA formation including matrix metalloproteinases (MMPs) and TIMP (tissue inhibitors of MMPs) within the aneurysm wall ^5^. In patients, hemodynamic parameters are mainly studied by computational fluid dynamics (CFD) calculated on high resolution data sets such as 3D digital subtraction angiography (DSA). Recently CFD on high-resolution 9.4T MRI data sets has been performed to image mouse circle of Willis ^6^. However, this high-field MRI is poorly available for preclinical research. Therefore, there is a need to develop a simple and reliable imaging technique to study IA formation and hemodynamic alteration in intracranial arteries in mice. In this study, we used transcranial Doppler (TCD) imaging to evaluate IA formation and development by monitoring arterial tortuosity and blood flow velocities in intracranial arteries in 2 well-known mouse models of IA.

## Materials and Methods

### Mice

C57BL6 mice were bred and housed in our local conventional animal facility. All experiments were performed in accordance with French ethical laws (agreement number: APAFIS n°3962). All animals were maintained under standard conditions with food and water given *ad libitum*.

### Intracranial aneurysm induction

Intracranial aneurysms were induced as described by Hashimoto *et al*., and Hoh *et al*. in 8-10 week-old C57BL6 mice ^7,8^.

Briefly, in Hoh’s model, chronic hypertension is induced *via* right renal artery and left common carotid artery ligations in deeply anesthetized female mice. One week later, stereotaxic injection (coordinates: 1,2 rostral; 0,7 lateral; 6 mm deep) of elastase (1U/mL diluted in 10μL of saline solution) was performed, followed by subcutaneous implantation of an osmotic pump (ALZET 2004) filled with angiotensin II (1000 ng/kg/min). In addition, β-aminopropionitrile was delivered in drinking water (0,12%, Sigma Aldrich) throughout the model (Fig. 1A).

**Figure 1:**
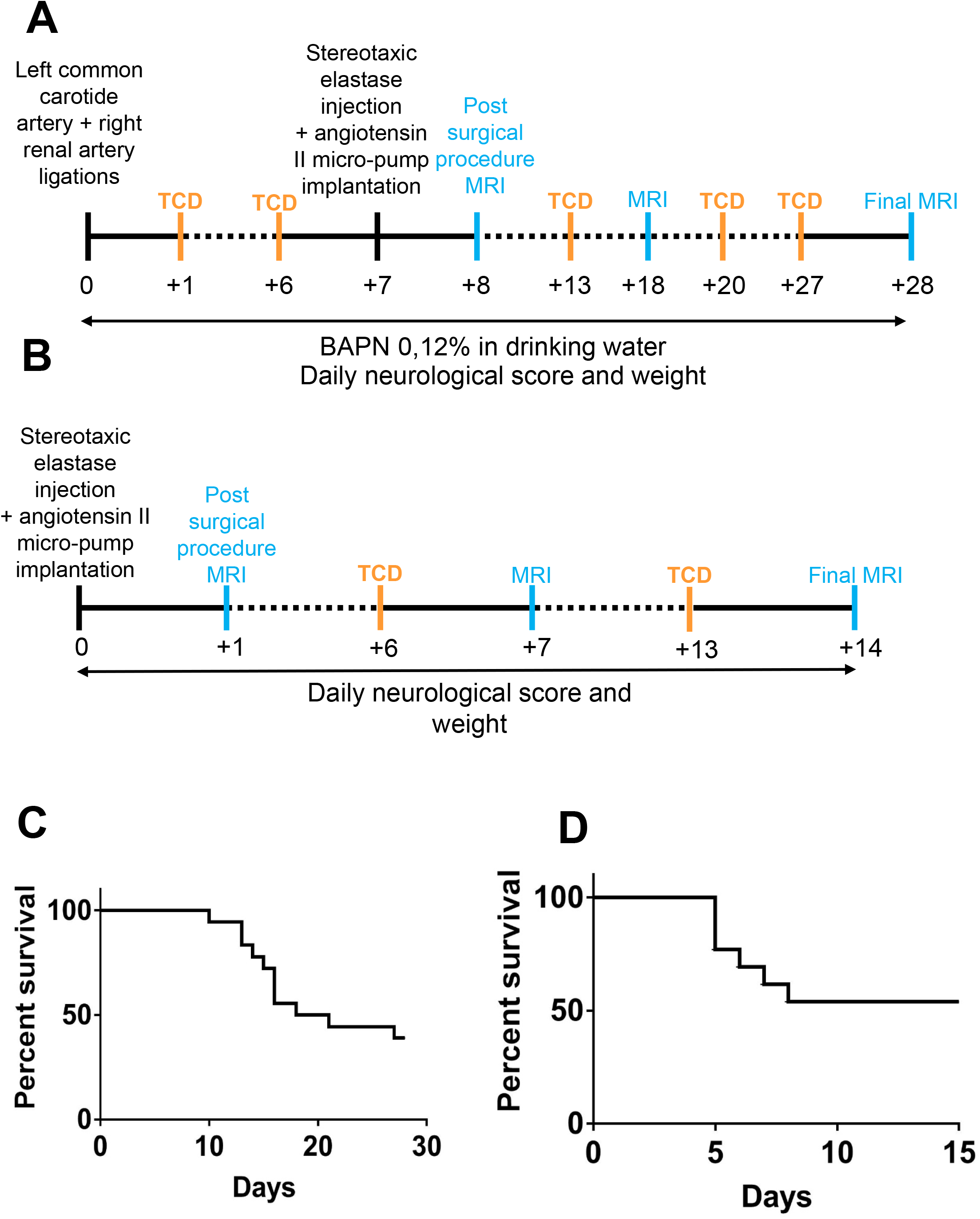
Schematic time-course of Hoh’s (**A**) and Hashimoto’s (**B**) models. TCD imaging and MRI are performed once a week and the surgical procedure is detailed in the materials and methods section. Survival curve of mice subjected to Hoh’s (**C**) or Hashimoto’s (**D**) model.

Hashimoto’s model consists in a stereotaxic injection in the right basal cistern (coordinates: 2,5 posterior; 2,5 lateral and 6 mm deep) of 2,5 μL of elastase (35 mU diluted in saline solution) in deeply anesthetized male animals, followed by subcutaneous implantation of an osmotic pump (ALZET 2004) filled with angiotensin II (600 ng/kg/min, Sigma Aldrich) (Fig. 1B).

For each model, a magnetic resonance imaging (MRI) was performed the day following elastase injection to exclude animals exhibiting an intracranial hemorrhage. Once a week, each animal was scanned by MRI and intracranial blood flow velocity (BFVel) was assessed by TCD. Animals were observed daily for weight loss or onset of neurologic.

### Magnetic Resonance Imaging (MRI)

MRI images were acquired on a Pharmascan 7T/12 cm system using surface coils (Brucker, Germany). During acquisitions, anesthesia of thermoregulated mice was maintained using isoflurane 1,5%. T2*-weighted sequences were acquired using a FLASH (fast low angle shot) sequence and used to detect intracranial hemorrhage in hyposignal. 3D T1-weighted sequence was performed to visualize IA as an abnormal dilation of intracranial arteries.

### Transcranial Ultrasound Doppler

Thermoregulated mice were subjected to ultrasound measurements under isoflurane anesthesia (1.0%) using an echocardiograph (ACUSON SEQUOIA, Siemens, Erlangen, Germany) equipped with a 14.5-MHz linear transducer. The transducer was placed at the back of the head and neck allowing a horizontal cross-sectional view of the base of the skull. The color-coded Doppler then drew the circle of Willis. The Doppler sample of the pulsed-Doppler was placed successively in the left (lICA) and right (rICA) internal carotid arteries in the left and right anterior (lACA, rACA) and posterior (lPCA, rPCA) cerebral arteries, then in the basilar trunk (BT) for Doppler velocity waveforms recordings. Spectral analysis of the Doppler velocity waveforms was recorded at the different time-points in each models of IA. Time-averaged-spatial-averaged mean blood flow velocities (mBFVels) were then measured after Doppler beam-longitudinal axis of the artery angle correction. Heart rates were recorded to ensure the absence of cardiorespiratory depression during mBFVels acquisitions. The mBFVel measurements were previously investigated in control mice at basal states for intra-observer repeatability. Two series of measurements separated by a 10 min interval were recorded in ten mice. The repeatability coefficient (RC) was calculated as defined by the British Standard Institution (British standards institution: precision of test method (BS5497, part I) 1979), i.e. according to the formula RC^2^=∑Di^2^/n, where Di is the relative (positive or negative) differences within each pair of measurements and n is the sample. The intra-observer-RC values were 0.3, 0.4, 0.4 and 0.4 cm/s for ICAs, ACA, PCA and BT, respectively, largely inferior to the differences exhibited between the different values recorded at the different time-points. Differences were then considered as significant.

### Histology

Each mouse was deeply anesthetized and transcardially perfused with saline (25mL) followed by 20 mL of fixative solution (0.5% zinc chloride, 0.5% zinc acetate in 0.1 M Tris base buffer containing 0.05% calcium acetate, pH 7.4). Brains were post-fixed with zinc fixative solution (48h, 4°C), paraffin embedded and microtome transversal sections (10 μm) were collected. The sections were stained with orcein to visualize arterial elastic lamina.

### Analysis

TCD images were graded in a blinded manner. This grade scale is based on apparent tortuosity of the circle of Willis arteries. Grade 0: normal appearing arteries compared to healthy animals (and retrograde blood flow in left ICA and ACA in Hoh’s model); Grade 1: slight tortuosity (1 curvature); Grade 2: pronounced tortuosity (3 curvatures), Grade 3: advanced tortuosity (4 curvatures and more). The 4 grades were identified in Hoh’s model, but only grades 0 and 1 were observed in Hashimoto’s model.

The wall shear stress (WSS) was calculated according to this formula: WSS = μ × ((8×mBFVel)/D) in dynes.cm^-2 9^, where μ is the dynamic blood viscosity (0,04 g.cm-1.s^-1^), mBFVel the mean blood flow velocity and D the diameter of the vessel ^10^.

### Statistical analysis

Results are represented as mean ± S.E.M. Data and statistical processing were performed with GraphPad Prism 7 software. Statistical significance across groups defined by one factor was performed using Kruskall-Wallis test followed by a nonparametric t-test (Mann-Whitney test). The level of statistical significance was set at p < 0.05.

## Results

To study the usefulness of TCD imaging in the detection of brain aneurysms, we employ two mouse models of cerebral aneurysm described in the literature, respectively referred as Hoh’s and Hashimoto’s model ^7,8^.

### Intracranial aneurysm imaging by TCD and MRI in Hoh’s mouse model

In control mice, an anterograde cerebral blood flow (in red on TCD images) originating from left and right ICA mainly supply the left and right ACA respectively (Fig. 2A). A retrograde blood flow (appearing in blue on TCD images) originating from both the ICA and the BT supply the bilateral PCAs (Fig. 2A). All these arteries analyzed on MRI images present a straight shape and a homogeneous diameter. Moreover, analysis of the histological staining shows an intense and continuous orcein staining of the elastic lamina.

**Figure 2:**
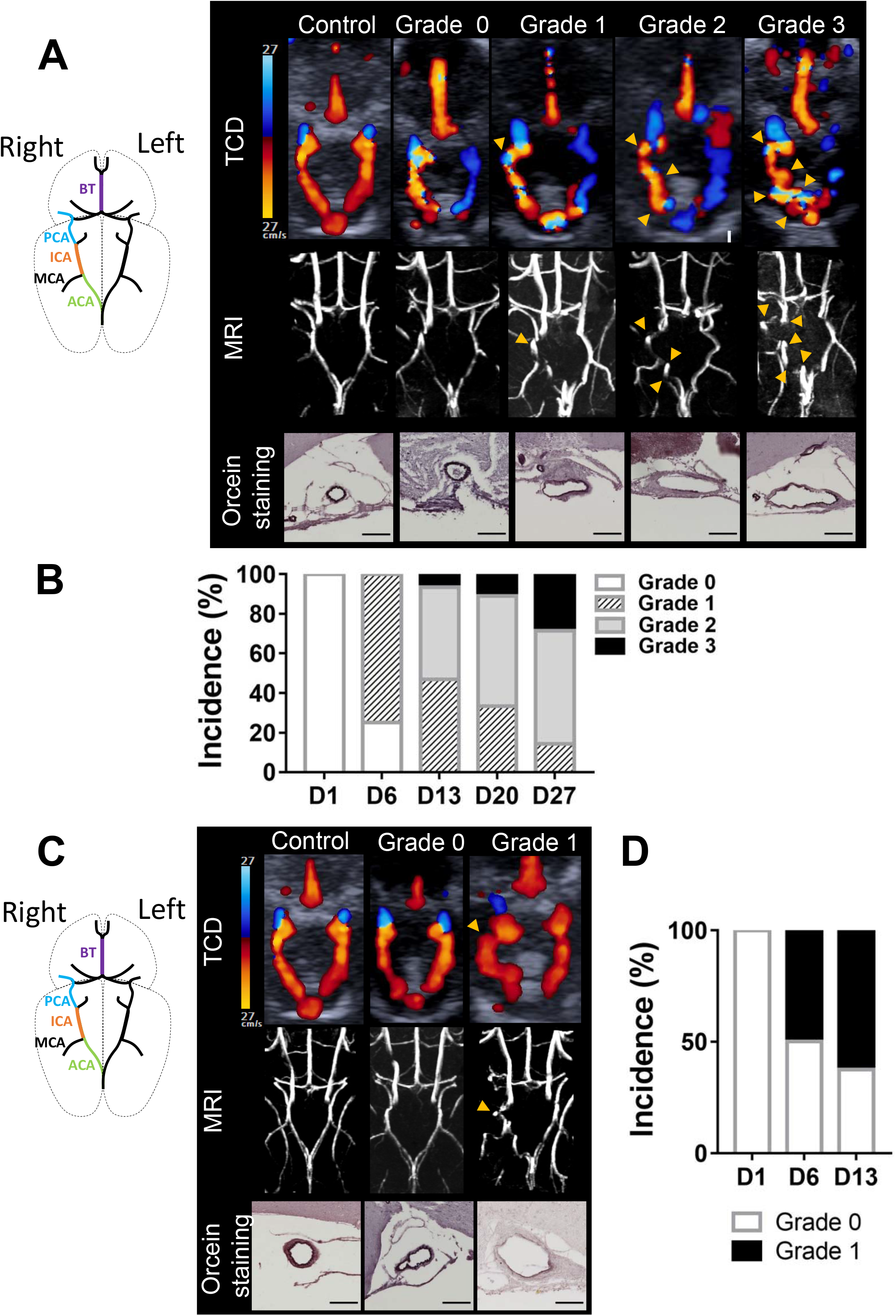
Increased arterial tortuosity detected by TCD imaging and confirmed by MRI uncover IA formation. Schematic representation of intracranial arteries. Representative TCD and T1w-MRI images of control, grade 0, 1, 2 and 3 mice subjected to Hoh’s model (**A**) and Hashimoto’s model (**C**) and corresponding representative orcein staining of the right ICA (deep purple: elastic lamina; scale bar: 100μm). Arrow-heads indicate abnormal curvatures. Percentage of apparent tortuous arteries, on TCD images, within the circle of Willis during the course of Hoh’s model (**B**) and Hashimoto’s model (**D**). BT: Basilar trunk; PCA: posterior cerebral artery; ICA: Internal carotid artery; MCA: Middle cerebral artery; ACA: anterior cerebral artery.

In mice subjected to Hoh’s model, one day after left common carotid artery ligation, we observed by TCD the rerouting of the cerebral blood flow in the circle of Willis. The left ICA and ACA were supplied *via* a retrograde blood flow originating from the anterior azygos cerebral artery (equivalent to the anterior communicant artery in human) (Fig. 2A, grade 0). This ligation associated with a stereotaxic elastase injection in hypertensive mice led to the development of intracranial arterial tortuosity on right ICA and ACA observed in both TCD and MRI (Fig. 2A, grade 1, 2 and 3, yellow arrows). Furthermore, this tortuosity was associated with IA formation characterized by elastic lamina disruption and degradation and artery dilation (Fig. 2A, grade 2 and 3). At day 1, all mice were scored as grade 0 and as soon as day 6, 75% of mice presented a grade 1 tortuosity (Fig. 2B). From day 13 to 27, mice presenting a grade 2 tortuosity increased from 46% to 57% and grade 3 tortuosity increased from 6% to 28% (Fig. 2B). At day 28, 76.9% of animals developed an IA, and 38% survived (Fig. 1C).

Due to the left ICA ligation, a marked increase of the mean blood flow velocity (mBFVel) in the right ICA was measured after the surgery in grade 0 compared to control mice, from 13.5 ± 0.8 to 22.4 ± 1.6 cm.s^-1^ (Fig. 3A). This mBFVel increases is maintained in grade 1 mice (24.1 ± 1.3 cm.s^-1^) but significantly decreases in grade 2 and 3 mice compared to grade 0 and 1 mice (18.6 ± 1.3 cm.s^-1^; Fig. 3A). Moreover, the correlation between the mBFVel and the right ICA diameter measured by MRI shows a significant correlation (R^2^: 0.405, p=0.0013; Fig. 3B). It can be noticed that mice with a high tortuosity grade (grade 2 and 3: orange dots) present a more severe artery dilation and mBFVel decrease than grade 1 (blue dots) and grade 0 mice (grey dots). Due to the increased mBFVel induced by the left common carotid artery ligation, the WSS significantly increases in the right ICA of grade 0 mice compared to control mice (control: 158.1 ± 8.4, grade 0: 241.5 ± 12.5 dynes.cm^-2^). However, we observed that the higher the mice were graded for their arterial tortuosity, the lower their calculated WSS were (grade 1: 169 ± 17.4, grade 2 and 3: 117.4 ± 9.7 dynes.cm^-2^; Fig. 3C).

**Figure 3:**
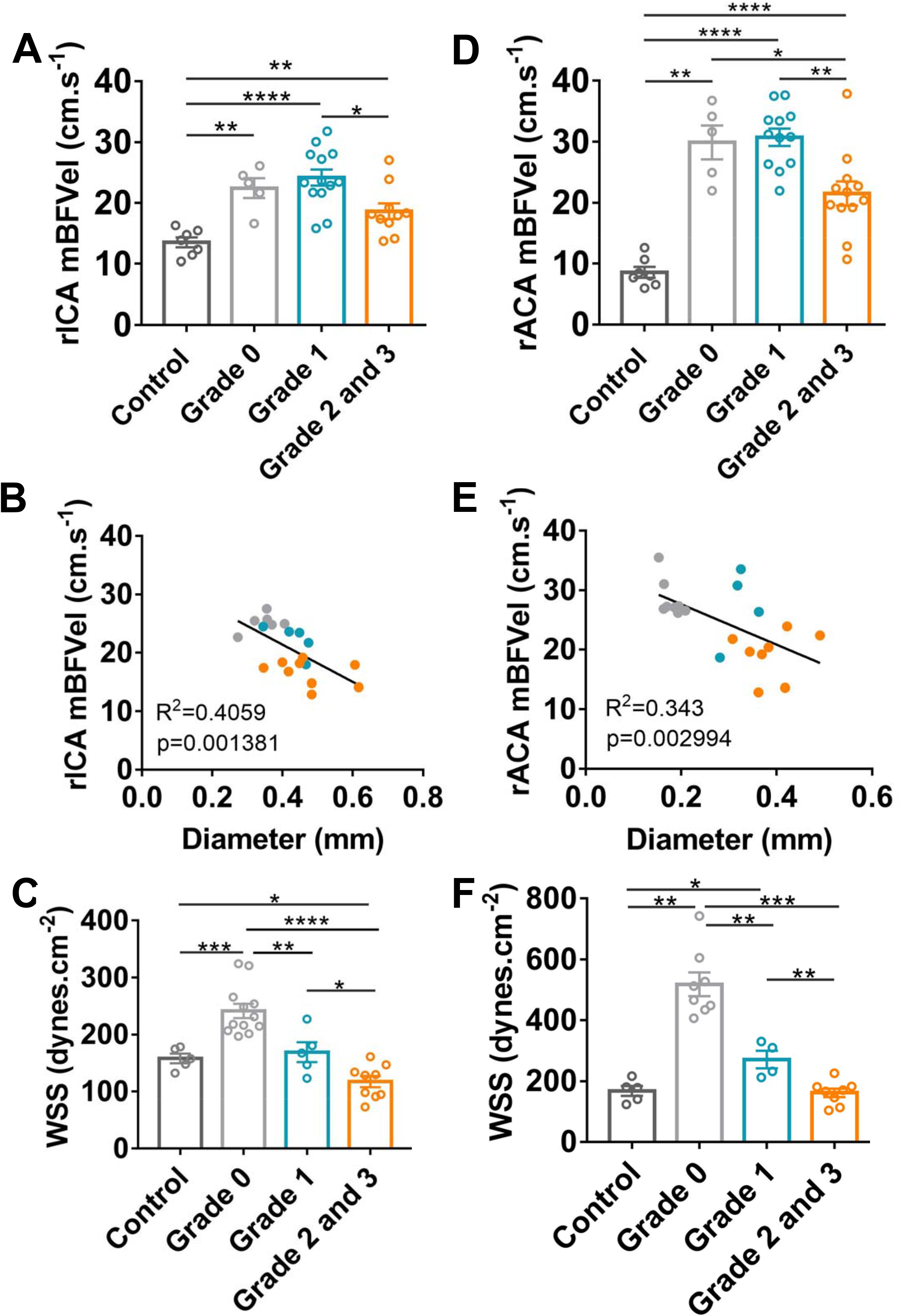
Hemodynamic parameters are altered in highly tortuous intracranial arteries in Hoh’s model. **(A-D)** Measurements of the mBFVel in the ICA or ACA by TCD of control and mice subjected to the Hoh’s model. **(B-E)** Correlation between the mBFVel and right ICA or ACA diameter measured by MRI (grey: grade 0; blue: grade 1; orange: grade 2/3). **(C-F)** Calculated WSS in the right ICA and ACA of control and mice subjected to Hoh’s model, according to their tortuosity grade. Data are represented as mean ± SEM; Mann-Whitney test; * p<0.05; ** p<0.01, *** p<0.001, and **** p<0.0001.

Similarly, in the right ACA where arterial tortuosity is observed (Fig. 2A, grade 2 and 3), the mBFVel is increased in grade 0 and 1 mice compared to control mice (control: 8.5 ± 0.8 cm.s^-1^ vs grade 0: 29.8 ± 2.7 cm.s^-1^ and grade 1: 30.7 ± 1.4 cm.s^-1^) and significantly decreases in grade 2 and 3 mice compared to lower grades (grade 2 and 3: 21.5 ± 1.9 cm.s^-1^; Fig. 3D). As for the right ICA, there is a significant correlation between the mBFVel and the right ACA diameter (R^2^: 0.343, P=0.0029; Fig. 3E), and the calculated WSS significantly decreases as the tortuosity grade increases (grade 0: 518 ± 38.9, grade 1: 271.2 ± 28.8, grade 2 and 3: 161.4 ± 14 dynes.cm^-2^; Fig. 3F).

### Intracranial aneurysm imaging by TCD and MRI in Hashimoto’s mouse model

In mice subjected to Hashimoto’s model, stereotaxic elastase injection associated to angiotensin-II induced hypertension led to the development of ICA tortuosity, revealed by TCD imaging and confirmed through MRI (Fig. 2C, yellow arrows). This apparent tortuosity is associated with altered vessel wall integrity characterized by elastic lamina degradation and arterial dilation (Fig. 2C, grade 1 orcein staining). From day 6 to 13, mice presenting a grade 1 tortuosity increased from 50% to 62,5% (Fig. 2D). At the end of this IA mice model, 66% of animals developed an IA and 53% survived (Fig. 1D).

In the right ICA, the mBFVel was significantly decreased in grade 0 mice compared to control mice (13.1 ± 0.8 vs. 18 ± 0.6 cm.s^-1^). The mBFVel is further reduced in grade 1 mice (presenting arterial tortuosity on TCD and MRI images) compared to control and grade 0 mice (grade 1: 8.9 ± 0.8 cm.s^-1^; Fig. 4A). In addition, there is a significant correlation between the right ICA mBFVel and its diameter, measured by MRI (R^2^: 0.472, p =0.006; Fig. 4B). As in Hoh’s model, we observed that mice with arterial tortuosity (grade 1, orange dots) present a more severe artery dilation and mBFVel decrease than grade 0 (blue dots) and control mice (grey dots) (Fig. 4B). Furthermore, the calculated WSS was significantly decreased in grade 0 and 1 mice compared to controls (grade 0: 110.6 ± 4.3 and grade 1: 83.5 ± 8.7 *vs*. control: 206.3 ± 16.7 dynes.cm^-2^) and was significantly more pronounced in grade 1 mice compared to grade 0 mice (Fig. 4C).

**Figure 4:**
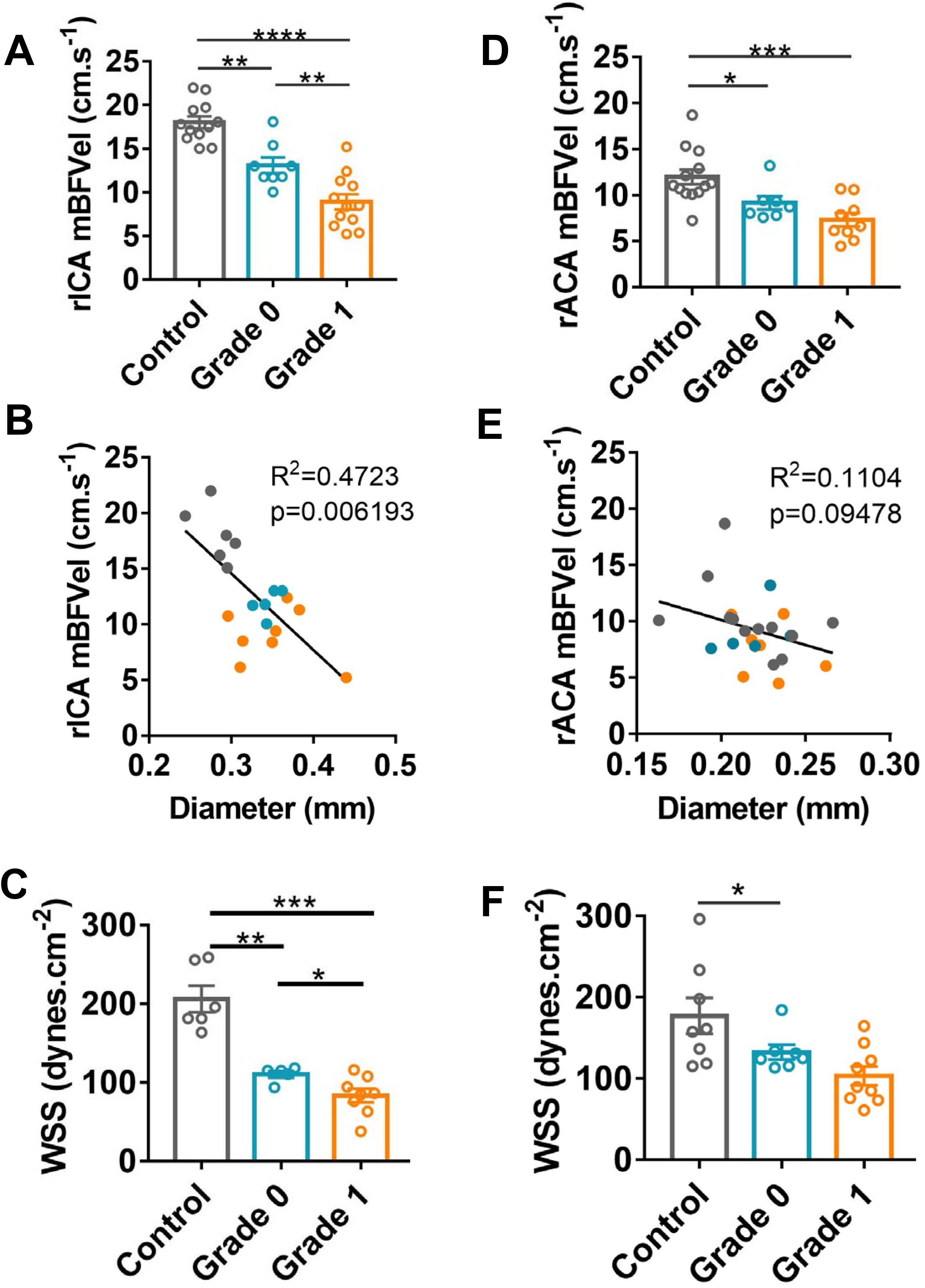
Hemodynamic parameters are altered in intracranial tortuous artery in Hashimoto’s model. (**A-D**) Comparison of the mBFVel, measured by TCD, in the right ICA and in the right ACA of control and Hashimoto’s model subjected mice, according to their tortuosity grade. **(B-E)** Correlation between mBFVel and right ICA or ACA diameter measured by MRI (grey: control; blue: grade 0; orange: grade 1). **(C-F)** Calculated WSS in the right ICA and ACA of control and Hashimoto’s model subjected mice, according to their tortuosity grade. Data are represented as mean ± SEM; Mann-Whitney test; * p<0.05; ** p<0.01, *** p<0.001, and **** p<0.0001.

A decrease of the mBFVel in the right ACA was also observed in grade 0 and grade 1 mice compared to control mice (Fig. 4D, grade 0: 9,1 ± 0.7 and grade 1: 7.3 ± 0.7 vs. control: 11.9 ± 0.8 cm.s^-1^). Interestingly, in agreement with the absence of apparent tortuosity on the right ACA on TCD and MRI images, no difference of mBFVel was observed between grade 0 and 1 mice (Fig. 4D). Confrontation of the MRI and TCD data showed no correlation between the mBFVel and the arterial diameter for the right ACA (R^2^: 0,110; p = 0,09; Fig 4E). In addition, there was no difference between the calculated WSS of grade 0 and grade 1 mice (respectively 132.8 ± 9 and 103.4 ± 11.6 dynes.cm^-2^; Fig. 4F).

## Discussion

The gold standard to image brain aneurysms in preclinical models is based on MRI followed by histological analysis to confirm IA formation in animal models. Those methods are expensive, time consuming and not readily available. In addition, it requires specific post-treatment imaging software to visualize the brain vasculature. Here, we use the ultrasound imaging as a novel method to longitudinally study intracranial aneurysm formation in mice.

There are several models of IA used in large animals (pigs, dogs, rabbits) or small animals (rodents). In mice, there are three main models: (i) the Morimoto’s model describing aneurysm formation at the site of anterior cerebral artery and olfactory artery bifurcation. The main limitations of this model are the time frame of the aneurysm formation (> 4 months) and their very small size (arterial wall protrusion of a few micrometer). (ii) The Hashimoto’s model, based on a stereotaxic elastase injection in hypertensive mice, allows the study of IA formation, (iii) the Hoh’s model is a combination of the two previous models including the ligation of the left common carotid and the right renal arteries prior to the elastase injection in hypertensive mice. This model allows the study of the aneurysm formation and rupture.

In Hashimoto’s and Hoh’s models, we uncover the presence of arterial tortuosity which is significantly correlated with the presence of an aneurysm confirmed by MRI and histology analysis. We also demonstrated that a high grade of tortuosity is associated with a significant decrease of the mean blood flow velocities in the internal carotid arteries and a greater arterial dilation. In line with our findings, patients with high ICA tortuosity present intracranial aneurysm ^11,12^. Moreover, patients with high arterial tortuosity display a lower shear stress which can promote matrix metalloproteinase activation and the weakening of the arterial wall ^13^ and leukocyte infiltration ^14^. Clinical and experimental studies strongly suggest that mechanical factors such as blood pressure, blood flow, axial tension and wall structural changes play an important role in the development of arterial tortuosity ^15^. In addition, computational studies have shown that lumen shear stress and wall stress are altered in tortuous arteries ^16,17^. Nevertheless, the molecular mechanisms in arterial tortuosity remain unknown.

In mice as well as in humans, alteration of the blood flow velocities plays a key role in the variation of the arterial diameter ^18^. Previous studies showed that an increase in blood velocities induces an outward physiological adaptation of the vascular wall ^19^. In the Morimoto model ^20^ induced by arterial ligation, small vascular protrusions characterized by some fragmentations of the internal elastic lamina are formed in several months which may reflect a slow remodeling process. To the contrary, in the Hashimoto and Hoh models ^7,8^, the arterial wall degradation induced by the elastase and hypertension contribute to the accelerated dilation and the formation of tortuous aneurysms highlighting a pathological outward remodeling rather than a growth of the arterial wall. Lower blood flow velocities and WSS, as observed in both models, have been shown to induce the expression of proinflammatory molecules ^21^ and the release of vasoactive molecules by endothelial cells and smooth muscle cells ^18,22^. The recruitment of leukocytes to the diseased artery and the vascular tone are key components in the outward remodeling ^8,23^.

Even though TCD appears to be a promising new imaging technique for IA in animal models, our study presents some limitations. Indeed, TCD imaging does not detect IA located on circle of Willis branches, such as the middle cerebral artery, despite the IA presence confirmed through MRI and histology. Moreover, arterial diameter cannot be accurately measured on TCD images as the spatial resolution of the TCD modifies the apparent artery diameter. Finally, the resolution of TCD imaging does not allow the detection of small IA, such as the ones developed in the Morimoto’s model.

In conclusion, our work shows, for the first-time, the advantage of TCD as a complementary method to image brain aneurysms in preclinical models. Our findings provide a rationale for assessing the tortuosity and the drop of the intracranial blood flow velocity to predict IA severity in mice.

## Author Contributions

HL, AB, CM, JS and PB performed experiments. HL conducted data analysis. HL, PB and YB designed the research, interpreted the data and wrote the paper.CO provided guidance in the set-up of the animals models. MM contributed to the interpretation of MRI images and provided feedback on animal models. All authors provided critical comments and edited the manuscript.

## Acknowledgements

We thank David S. Paul for editing the manuscript.

